# Beyond Pathway Boundaries: A Degree-Aware Network Clustering Test for Gene Sets

**DOI:** 10.64898/2026.05.04.722765

**Authors:** Bryan Queme, Paul Marjoram, Huaiyu Mi

## Abstract

Over-representation analysis (ORA) is the most commonly used interpretation tool for gene lists despite well-documented limitations: pathway boundaries are fixed, genes are assumed independent, and results depend on the background set. Network-based methods address these using interaction-network modularity, but introduce hub bias: highly connected genes appear clustered under naive nulls because curated networks overrepresent well-studied genes. Existing corrections are imperfect: edge permutation destroys the topology the test should condition on, and propagation methods hide the confound in parameter tuning. We introduce MANGO (Moran’s Autocorrelation for Network Gene Over-representation), which asks one conditional question: does a gene set’s spatial autocorrelation on a fixed biological network exceed what its degree composition alone would predict? MANGO computes Global Moran’s I under a null that conditions on both the network and the binned degree distribution of the gene set, then decomposes significant signals at the component and gene level. In benchmarks, uniform nulls produce a false positive rate of 1.0 on hub-enriched gene sets with no real clustering; ten-bin degree-stratified nulls bring that to 0.0 with no power loss (AUC ≥ 0.98; on degree-typical signals, |ΔAUC| ≤ 0.004). Pathway-spiking simulations confirm detection of real biological clustering across diverse pathway sizes and degree profiles. Applied to the FIGI colorectal cancer GWAS (204 SNPs), the set is degree-typical (KS p = 0.83), yet Moran’s I is highly significant (p < 0.001). Component-level jackknife localizes the entire signal to a single 24-gene module spanning TGF-β, Wnt/cadherin, and related pathways, with four bottlenecks (SMAD3, MYC, CTNNB1, PTPN1) matching established CRC driver biology.

**eTOC blurb:** MANGO tests whether a gene set’s spatial autocorrelation on a biological network exceeds what its degree composition predicts, by conditioning Global Moran’s I on the binned degree distribution with the network held fixed. Significant signals are decomposed to modules, bottleneck genes, and statistical drivers through component jackknife, articulation-point, and gene-jackknife analysis.

## Introduction

Over-representation analysis (ORA) is the default interpretation tool for understanding the underlying biology for high-throughput methods with gene lists across computational biology, from GWAS-derived gene sets to differential expression results, knockout screens, proteomics hits, and any other workflow that produces a set of candidate genes^1^. The standard pipeline asks whether the implicated genes are over-represented^2^ in curated databases from Reactome^3^, KEGG^4^, PANTHER^5^, or the Gene Ontology^6,7^ using a Fisher’s exact or hypergeometric test^1^. ORA has been the field standard for twenty years and remains the default first pass in most downstream analyses, yet its limitations are thoroughly catalogued. Pathway boundaries are manually and arbitrarily defined, and do not reflect disease-specific connectedness, gene independence is assumed despite pervasive functional correlation, and results are sensitive to the database choice. Additionally, ORA tests each pathway independently, so cross-pathway biology is invisible to the test^8–10^.

Network-based methods address these limitations by evaluating whether input genes cluster on a molecular interaction network, exploiting the disease-module principle^11^. These methods fall into several families. Network propagation approaches such as HotNet2^12^ spread evidence from seed genes through the network via diffusion or random walks to identify significantly “hot” subnetworks. Dense-module searches such as dmGWAS^13^ expand modules around low-pvalue seeds to find connected subgraphs enriched for association signal. Active-module identification tools such as DOMINO^14^ partition the network into modules and test each for enrichment against permuted input. Neighborhood-based approaches such as LEAN^15^ test whether a gene’s direct network neighbors carry stronger association signal than expected. networkGWAS^16^ uses kernel regression on PPI neighborhoods within a mixed-model SNP-set framework. These methods are compared in Table 1. In principle, these methods sidestep pathway boundaries and exploit cross-pathway modularity. However, in practice, they inherit a new confound: curated interaction databases reflect both genuine biological connectivity and the tendency for well-studied genes to accumulate more annotated interactions^17,18^. Because these two sources of high degree are entangled, a method that does not control for degree cannot distinguish clustering driven by biological modularity from clustering driven by the degree distribution of the gene set. It’s been shown that node degree alone can account for roughly half of apparent guilt-by-association performance^19^, and DOMINO demonstrated that active-module algorithms report GO-enriched modules even on randomly permutated input^20^. A method that fails to correct for degree bias risks reporting clustering that is a property of the network rather than of the gene set.

**Table 1.**
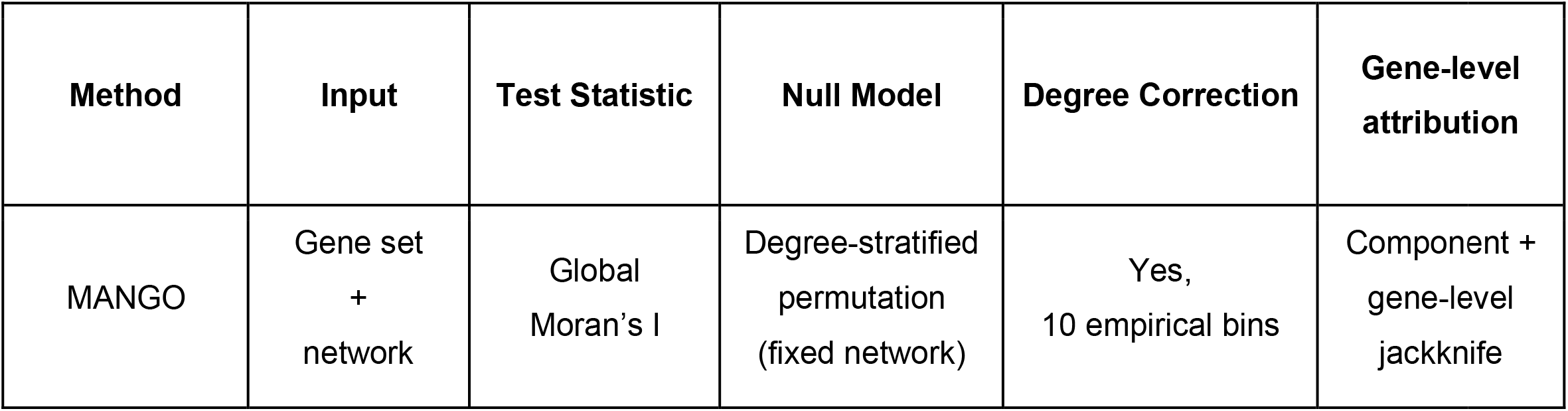
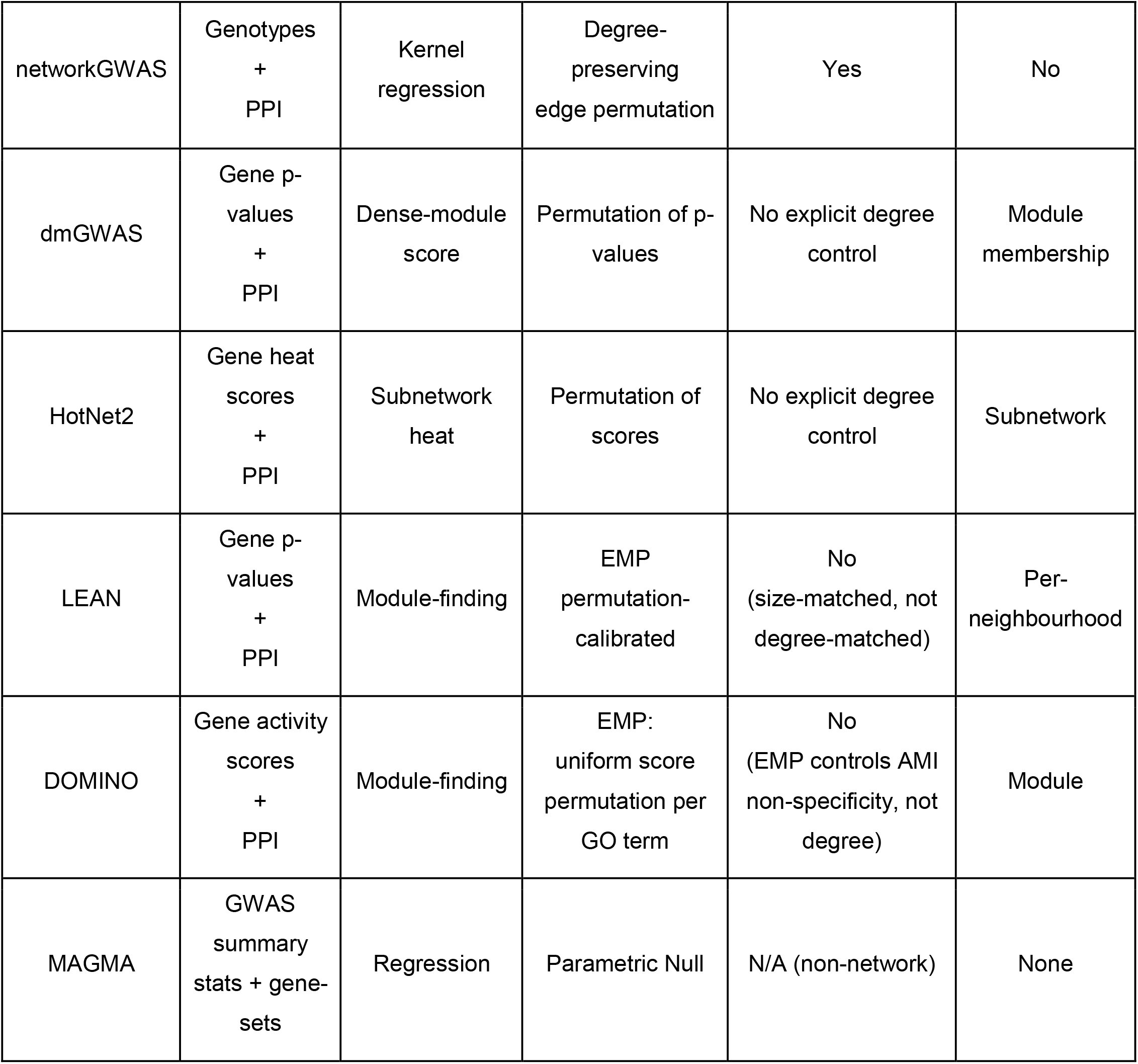
Comparison of MANGO with representative existing methods for network-based gene-set analysis.

| Method | Input | Test Statistic | Null Model | Degree Correction | Gene-level attribution |
| --- | --- | --- | --- | --- | --- |
| MANGO | Gene set + network | Global Moran's I | Degree-stratified permutation (fixed network) | Yes, 10 empirical bins | Component + gene-level jackknife |
| networkGWAS | Genotypes<br>+<br>PPI | Kernel<br>regression | Degree-preserving<br>edge permutation | Yes | No |
| dmGWAS | Gene p-<br>values<br>+<br>PPI | Dense-module<br>score | Permutation of p-<br>values | No explicit degree<br>control | Module<br>membership |
| HotNet2 | Gene heat<br>scores<br>+<br>PPI | Subnetwork<br>heat | Permutation of<br>scores | No explicit degree<br>control | Subnetwork |
| LEAN | Gene p-<br>values<br>+<br>PPI | Module-finding | EMP<br>permutation-<br>calibrated | No<br>(size-matched, not<br>degree-matched) | Per-<br>neighbourhood |
| DOMINO | Gene activity<br>scores<br>+<br>PPI | Module-finding | EMP:<br>uniform score<br>permutation per<br>GO term | No<br>(EMP controls AMI<br>non-specificity, not<br>degree) | Module |
| MAGMA | GWAS<br>summary<br>stats + gene-<br>sets | Regression | Parametric Null | N/A (non-network) | None |

Several approaches to degree correction have been proposed, but each holds different aspects of the data fixed under the null, with consequences for interpretation. The configuration model^21^ rewires edges while preserving degree, but destroys the topology the test should condition on. A question that recognizes that gene-set clustering on a network is a spatial autocorrelation should be: are labelled nodes (genes in the set) more adjacent to each other than expected? Global Moran’s I^22,23^ is the classical measure of spatial autocorrelation, quantifying whether neighboring observations carry similar values. A recent framework for Moran-type statistics on networks^24^ proposes both a uniform data-permutation null (which does not control for degree) and a configuration-model null (which randomizes the network). networkGWAS takes a different approach, combining circular SNP permutation with degree-preserving network permutation, but requires individual-level genotype data, limiting it to GWAS with full genotypes.

We propose a different strategy, which we call MANGO (Moran’s Autocorrelation for Network Gene Over-representation). Rather than rewire the network or permute gene scores uniformly, we hold the network fixed and ask one conditional question: given a gene set with a particular connectivity profile on this network, is its spatial autocorrelation stronger than what a degree-matched random draw would produce? The null is constructed by partitioning all network nodes into bins based the network degree distribution quantiles, then drawing a replacement gene set that preserved the number of genes in each bin. This ensures that a gene set enriched for highly connected genes is compared against other sets with the same proportion of highly connected genes, rather than against the network at large. The test statistic is Global Moran’s I, and the p-value is obtained through a permutation test. Moran’s I has been applied in spatial transcriptomics^25^ but not previously to gene sets on integrated biological networks with calibrated degree control. If the global test is significant, a second tier decomposes the signal: a component-level jackknife localizes which modules carry the signal, and a gene-level jackknife identifies which individual genes contribute to the statistic.

Here, we show why degree control matters for network-based gene-set analysis, demonstrate through simulation that uniform permutation nulls are miscalibrated on hub-enriched gene sets while degree-stratified permutation restores nominal error rates with negligible power cost, empirically determine the number of degree bins needed for reliable calibration, validate detection on real biological pathways across a range of sizes and degree profiles, and apply the framework to the FIGI colorectal cancer GWAS, where MANGO identifies a 24-gene module spanning TGF-beta, Wnt signaling, Cadherin, Gonadotropin, and Alzheimer’s pathway consistent with established CRC biology.

## Methods

### Biological Causal Network

We constructed an integrated gene-gene causal network from the PathwayCommons^26^ v12 BioPax^27^, restricting to three curated sources: Reactome, PANTHER, and KEGG. BioPAX interaction records were converted to a simple interaction format representing gene-level edges. We carefully analyzed PathwayCommons 10 interaction types and only kept the 4 that seemed causal (controls-state-change-of, controls-transport-of, controls-phosphorylation-of, and controls-expression-of) to reflect causal relationship between genes.

### MANGO test

Given a gene set S on a fixed biological network *G* = (*V, E*), does S cluster on G more than its connectivity profile, degree profile, alone would predict? MANGO answers this by computing a single conditional probability:

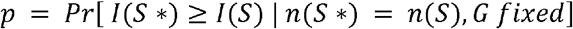

where I is Global Moran’s I, *n* (*S*) = (|*S* ⋂ *B*_1_|,..., |*S* ⋂ *B*_*B*_|) is the degree-bin signature of S, and *B*_*b*_ = {*v* ∈ *V*: *q*_*b*-l_ < *d*_*v*_ ≤ *q*_*b*_} are B bings defined by the empirical degree-quantiles of G. If p is small, the gene set cluster is beyond what a degree-match random draw would produce on the real network. Everything below operationalizes this statement.

#### Test Statistic

Global Moran’s I measures the spatial autocorrelation of a variable over a weighted graph, that is, the extent to which neighboring nodes carry similar values. Let *x* ∈ {0,1}^|*V*|^ be the gene-set indicator (*x*_*i* = l_ if gene i ∈ *S*, 0 otherwise), and let *w*_*ij*_ =1 if nodes *i* and *j* share an edge in *G*:

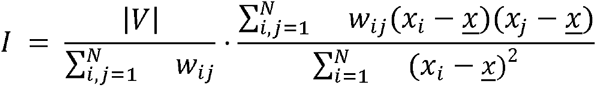

*I* > *E*[*I*] indicates that genes in *S* are adjacent to one another more often than expected. Binary adjacency is used throughout: two genes are “neighbours” if and only if the curated interaction network asserts a direct interaction between them.

#### Null Model

The probability in the equation above is estimated by permutation. All nodes in *G* are assigned to one of *B* = 10 bins by empirical degree quantiles of G (*B* = 1 is a uniform null with no degree control, *B* = 2 splits at the median, *B* = 3 at terciles, and so on). For each of P permutations, a null gene set *S*\* is drawn by sampling |*S* ∩ *B*_*b*_| genes uniformly without replacement from each bin *B*_*b*_ independently across bins. The network G is held fixed; only gene-set membership is permutated. *I*(*S* *) is computed on the same adjacency matrix, and the p-value is the fraction of null statistics meeting or exceeding *I*(*S*).

This null conditions on the network and on the connectivity profile of the gene set, which is what separates MANGO from a uniform permutation (*B* = 1, which does not control for degree) and from a configuration-model null (which randomises the network itself). The key design question is how many bins are needed. Too few bins leave residual degree confounding; too many shrink the per-bin candidate pool and can make the null overly conservative. We tested *B* = 1 through 10 across all simulation scenario (described below) to empirically determine the bin count at which false positive control stabilises without sacrificing power. The permutation is *P* = 1000 for the primary analysis.

#### Alternative statistics

Join Count, a classical spatial statistic that counts edges connecting labelled nodes (here, genes in the test set), was evaluated as an alternative to Global Moran’s I under the same degree-stratified null framework across all tested values of B. A fragmentation index, defined as the normalized sum of reciprocal component sizes in the induced subgraph (lower values indicate more genes concentrated in large components), was also tested as a measure of subgraph cohesion related to reciprocal-based fragmentation measured used in network robustness analysis^28^. The simulation results determine which statistics achieves the best combination of false-positive control and power. Full comparison including all statistics are in Supplementary Figures [].

### Post-hoc Decomposition

If the global test rejects, the clustering signal is real but unattributed. The second tier of MANGO decomposes the signal at two levels, analogous to moving from an omnibus F-test to post hoc contrasts in an ANOVA. The induced subgraph of S on G is the substrate for both.

We use the jackknife (leave-one-out recomputation) for attribution because it is compatible with the permutation framework: it is non-parametric, requires no distributional assumptions about gene-level contribution to Moran’s I, and measures each element’s marginal contribution to a single already-significant global statistic rather than testing each gene against its own null, which would introduce a severe multiple-testing burden. At the component level, the jackknife re-runs the full permutation test (yielding a proper p-value for the reduced set), while at the gene level it re-computes only the observed *I* (yielding a faster relative ranking without additional permutation).

#### Component jackknife (where does the signal live?)

The induced subgraph is partitioned into connected components. For each non-trivial component *C*, the genes of *C* are removed from the test set and Global Moran’s I and its degree-matched p-value are recomputed, yielding *p*_-*c*_. A larger increase Δ*p*_*c*_ = *p*_-*c*_ − *p*_*full*_ means component *C* carries a substantial fraction of the clustering signal. This is the coarsest and most powerful attribution step: it answers “which module matters?” before any gene-level analysis.

#### Gene-level jackknife (which gene drive the statistic?)

For each gene *i* in the test set, Global Moran’s I is recomputed with gene *i* removed, yielding *I*_*-i*_. The jackknife contribution Δ*I*_*i*_ =*I* - *I*_-*i*_ identifies statistical drivers: genes whose presence explains much of why Moran’s I is elevated, whether or not they are structurally critical.

Within each signal-carrying component, we also report articulation points (nodes whose removal fragment the component into two or more disconnected pieces, identified by Tarjan’s algorithm) as structural context for interpreting the jackknife results. Articulation points are a deterministic graph property rather than a statistical test, but they add interpretive value by distinguishing genes that serve as topological bridges between otherwise-separate parts of a module from genes that contribute edge density without holding the module together.

A schematic is shown in Figure 1.

**Figure 1.**
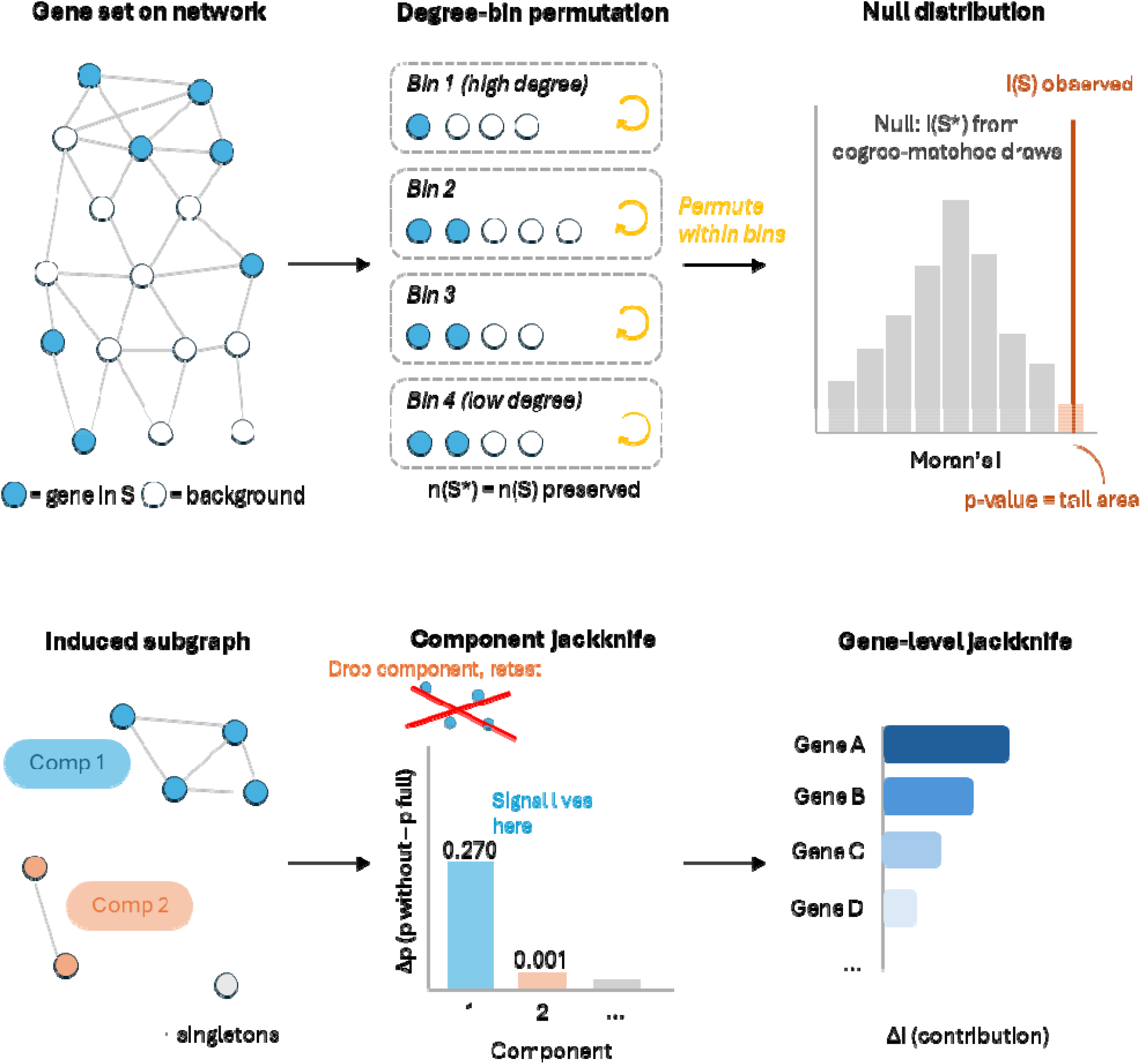
Overview of the MANGO framework. (a) Tier 1: a gene set is mapped onto the fixed biological interaction network; Global Moran’s I is computed, and its p-value is estimated by degree-stratified permutation that preserves the gene set’s connectivity profile. (b) Tier 2: if significant, the induced subgraph is decomposed via component-level jackknife (which module carries the signal) and gene-level jackknife (which genes drive the statistic), with articulation points reported as structural context.

### Simulation Framework

We benchmarked MANGO against a set of synthetic scenarios on the integrated network, each designed to test a specific calibration or power property. Signal and null gene sets were varied at n = 100, 200, and 500 total genes, with 100 replicates per signal-fraction level and 1,000 permutations per test.

#### Synthetic scenarios

In these scenarios, clustered signal genes are planted using breadth-first search (BFS): a seed node is chosen, and the cluster is grown outwards by successively adding all neighbors at each step until the target size is reached. This produces a connected, topologically compact cluster by construction, giving the test statistic a clear signal to detect. Because BFS preferentially visits high-degree nodes, the resulting clusters tend to be hub-enriched; scenario S7 explicitly controls for this by resampling BFS clusters until the Kolmogorov-Smirnov test confirms that the cluster’s degree distribution is not significantly different from the network background.

- **S1 (single tight cluster)**. 50 signal genes planted as one BFS-grown cluster, diluted with random off-pathway genes. Tests the detection of concentrated topological signal.
- **S2 (dispersed clusters)**. Three BFS clusters of 20 genes each, separated by at least 5 network hops. Tests the detection of a multi-module signal.
- **S3 (hub-enriched, no real clustering, the key false-positive control)**. 50 high-degree genes drawn from the full network but topologically dispersed, so there is no real clustering. A well-calibrated test must report this as non-significant under the degree-matched null, even though the hubs would create apparent clustering under a uniform null.
- **S3b (degree-typical null, calibration control)**. 50-degree-typical genes (KS test p > 0.05 against the background degree distribution) drawn at random. No degree confound, no real clustering. Tests whether the degree-matched null is nominally calibrated or over-conservative.
- **S7 (degree-matched BFS cluster, power-cost assessment)**. 50 signal genes from a BFS cluster whose degree distribution approximates the network background (not significantly different by KS test). Signal fraction was systematically varied (5%, 10%, 20%, 40%, 60% of the total gene set) across set sizes K = 100, 200, and 500 to assess whether degree-matched nulls sacrifice statistical power as the proportion of clustered signal genes changes. This is the primary test of whether degree control is free when the signal is degree-typical.
- **S8 (degree-typical dispersed signal, realistic GWAS-like case)**. A mixture of small BFS clusters and isolated singletons, all with background-typical degree. Tests detection of mild, realistic clustering.

For each scenario, MANGO was run with a uniform (unstratified) permutation null and with degree-stratified nulls at 1–10 bins. We report the false positive rate at α = 0.05 from 200 replicates on S3 and S3b, and AUC from 100 replicates per signal level on S1, S2, S7, and S8. The primary real-data analysis used 1,000 permutations.

#### Pathway-spiking scenarios

The synthetic scenarios above plant signals via BFS walks, which produce connected clusters, but do not necessarily test whether MANGO detects the known biological clustering that actually exists in curated biological pathways. We therefore ran two additional simulation tiers using real pathway gene sets from the integrated network as the signal.

In the single-pathway tier, 8 pathways in the integrated network were filtered from those forming a single connected component in their induced subgraph, then stratified by gene count into three size classes (small: 20–50 genes; medium: 80–200; large: 300+). Within each size class, candidates were ranked by mean subgraph degree and two to three pathways were selected to represent a range of connectivity: small (HRR, 48 genes, mean degree 73; COPI transport, 44 genes, mean degree 28), medium (Clathrin-mediated endocytosis, 170 genes, mean degree 143; Smooth Muscle Contraction, 180 genes, mean degree 134), large (Olfactory Signaling, 435 genes, mean degree 23; Downstream TCR signaling, 475 genes, mean degree 174; BCR signaling, 522 genes, mean degree 192; FCERI-mediated Ca2+ mobilization, 500 genes, mean degree 190). Pathways are categorized by mean network degree into high (> 150), mid (100–150), and low (< 50) degree strata; the network-wide mean degree is 34.5. For each pathway, signal genes were the pathway members themselves; the gene set was diluted with random non-pathway genes to total sizes of 100, 200, and 500 (12 feasible of 24 combinations, since small pathways cannot fill large sets). Signal fraction (the proportion of the gene set composed of real pathway genes) was varied at 100%, 80%, 60%, 40%, and 20% by replacing pathway members with random genes.

In the multi-pathway tier, all single-component pathways with at least 30 genes were stratified by mean network degree into three pools: high (> 100), mid (50–100), and low (< 50). From these pools, seven sets of three pathways each were assembled to span the degree range. High-degree sets (mean network degree > 100): Set 1 (TCR / Insulin / Cadherin), Set 2 (HRR / BCR / Clathrin), Set 3 (VEGFR2 / Ca2+ / FCERI), Set 4 (SCF-KIT / DAP12 / Phagocytosis). Mid-degree: Set 5 (Chromatids / Rho / Inflammation). Low-degree: Set 6 (Olfactory / Neddylation / mTOR-AMPK). Set 7 is a synthetic degree-typical negative control (500 randomly selected genes whose degree distribution matches the network background by KS test, containing no real pathway clustering). Signal fraction was varied from 5% to 60% of the total gene set across K = 100, 200, and 500. This tests whether MANGO detects clustering from combinations of real pathways at varying dilution levels, and whether the degree-control cost depends on the degree profile of the pathways involved.

### Software and Reproducibility

Networks are represented as iGraph objects. Degree binning uses stats::quantile with ties broken by rank. Permutation testing is parallelized over cores. Code and the integrated network object are distributed at https://github.com/quemeb/MANGO

## RESULTS

### Biological Causal Network

The resulting undirected network contains 7,085 nodes and 122,219 edges, with a mean degree of 34.5 and a right-skewed distribution, typical of curated biological networks. The full degree distribution is shown in Supplementary Figure S1.

### Simulations: selecting the bin count and test statistic

#### Bin-count calibration

Under S3 (50 dispersed hub genes with no topological clustering, n = 200), Global Moran’s I with a uniform null (*B* = 1) has a false positive rate (FPR) of 1.000 at α = 0.05. That is, every replicate of a signal-free hub-enriched gene set is called significant. As *B* increases, the FPR drops steeply: 1.000 at B=2, 0.875 at B=4, 0.500 at B=5, 0.100 at B=6, 0.005 at B=7, and 0.000 from B=8 onward (Figure 2). The elbow is at 7-8 bins. We recommend B=10 as the default, a safe margin past the elbow that provides robust false-positive control without excessive fragmentation of the per-bin candidate pool.

**Figure 2.**
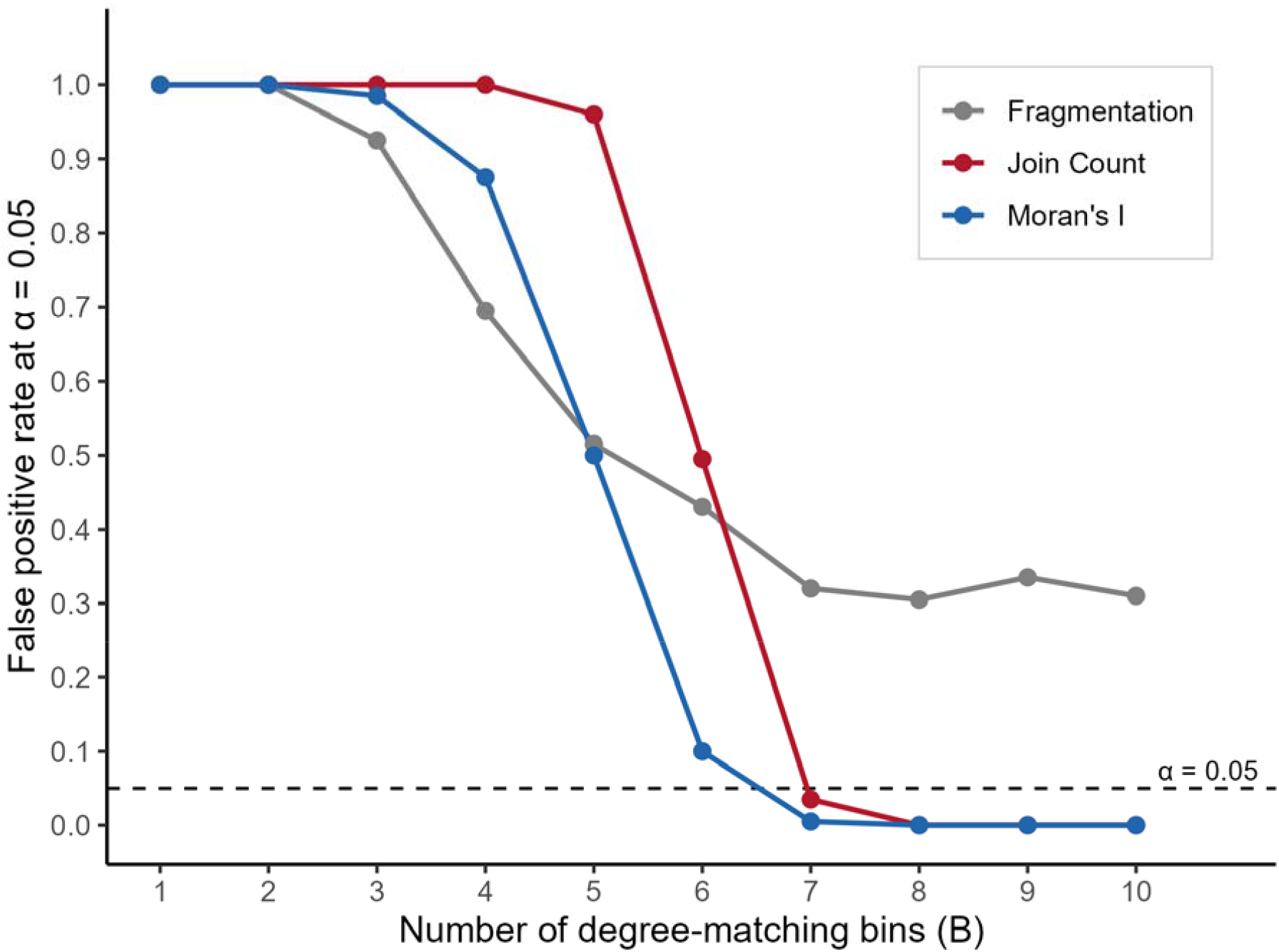
FPR calibration across bin count. False positive rate at α = 0.05 under the hub-enriched null (S3, n = 200) for Global Moran’s I, Join Count, and the fragmentation index as a function of the number of degree-stratification bins (B = 1– 10). Dashed line marks nominal α = 0.05.

#### Statistic selection

Join Count was tested under the same degree-stratified null across all values of. It achieves FPR control comparable to Moran’s I on hub-enriched nulls, but has lower power on dispersed-signal scenarios. The fragmentation index did not achieve nominal FPR at any bin count tested (FPR ≈ 0.31 on S3 at B=10, n = 200), likely because component-size distributions are less sensitive to degree stratification than spatial autocorrelation. Global Moran’s I with B=10 achieves near-zero FPR on hub-enriched nulls while retaining the best power profile across scenarios, and is therefore the recommended primary statistic.

#### Power

The B=10 null does not sacrifice power. On S1 (single tight cluster), AUC = 0.99 at n = 100 and 0.99 at n = 200, with TPR@FPR=0.05 of 0.90 at n = 200. On S2 (dispersed clusters), AUC = 1.00 at n = 100 and 0.98 at n = 200. On S7 (degree-matched BFS cluster with varied signal fraction), the result is strongest: across the full 5 × 3 grid of signal fractions (5%-60%) and set sizes (K = 100, 200, 500), AUC remained ≥ 0.98, and the mean AUC difference between uniform and nulls was −0.004 (range −0.031 to +0.011). Degree control is essentially free when the signal genes are degree-typical, regardless of how diluted the signal is. On S3b (degree-typical null, calibration control), the null has FPR of 0.010 at n = 200 and 0.035 at n = 500, close to the nominal 0.05, confirming that the procedure is calibrated rather than over-conservative (Figure 3).

**Figure 3.**
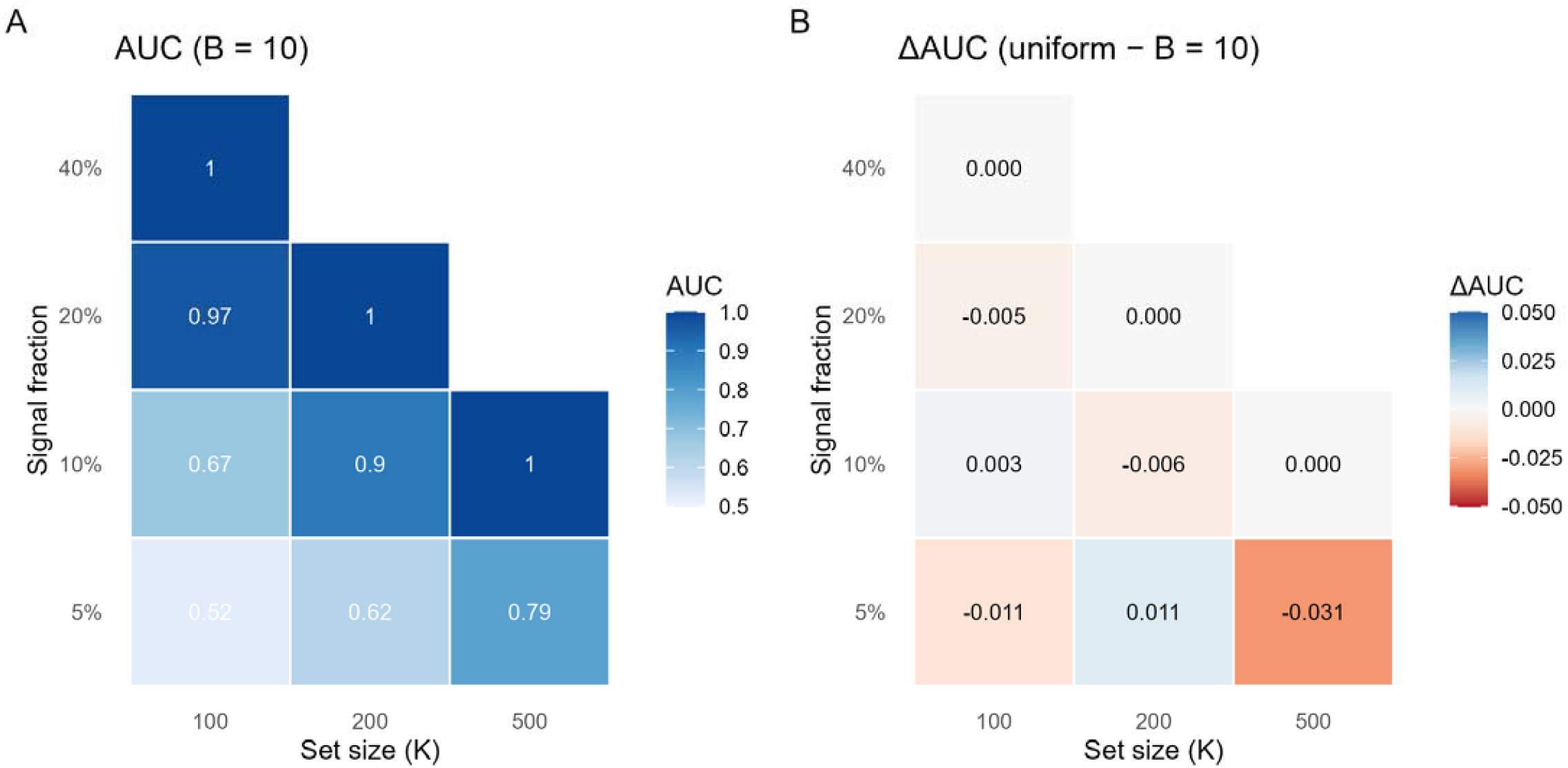
Degree control is free. (a) AUC of Global Moran’s I under B = 10 degree-stratified permutation across signal fractions and set sizes (S7). (b) ΔAUC (uniform minus B = 10), showing negligible power cost.

These findings establish Global Moran’s I with *B* = 10 degree-stratified bins as the primary MANGO detection test.

### Pathway-spiking simulations: MANGO detects real biological clustering

The pathway-spiking simulations test whether MANGO detects the clustering that exists in actual curated pathways rather than synthetic BFS clusters. In the single-pathway tier, high-degree pathways (TCR, BCR, FCERI Ca2+, Clathrin endocytosis) were detected with AUC ≥ 0.95 at moderate signal fractions across feasible set sizes. Lower-degree pathways (Olfactory Signaling, with mean network degree 22.9 compared to the network-wide mean of 34.5) were harder to detect, with AUC rising above 0.80 only at higher signal fractions or larger set sizes. This is expected: pathways whose genes are sparsely connected in the integrated network produce weaker spatial autocorrelation and require higher signal fractions to detect (Figure 4).

**Figure 4.**
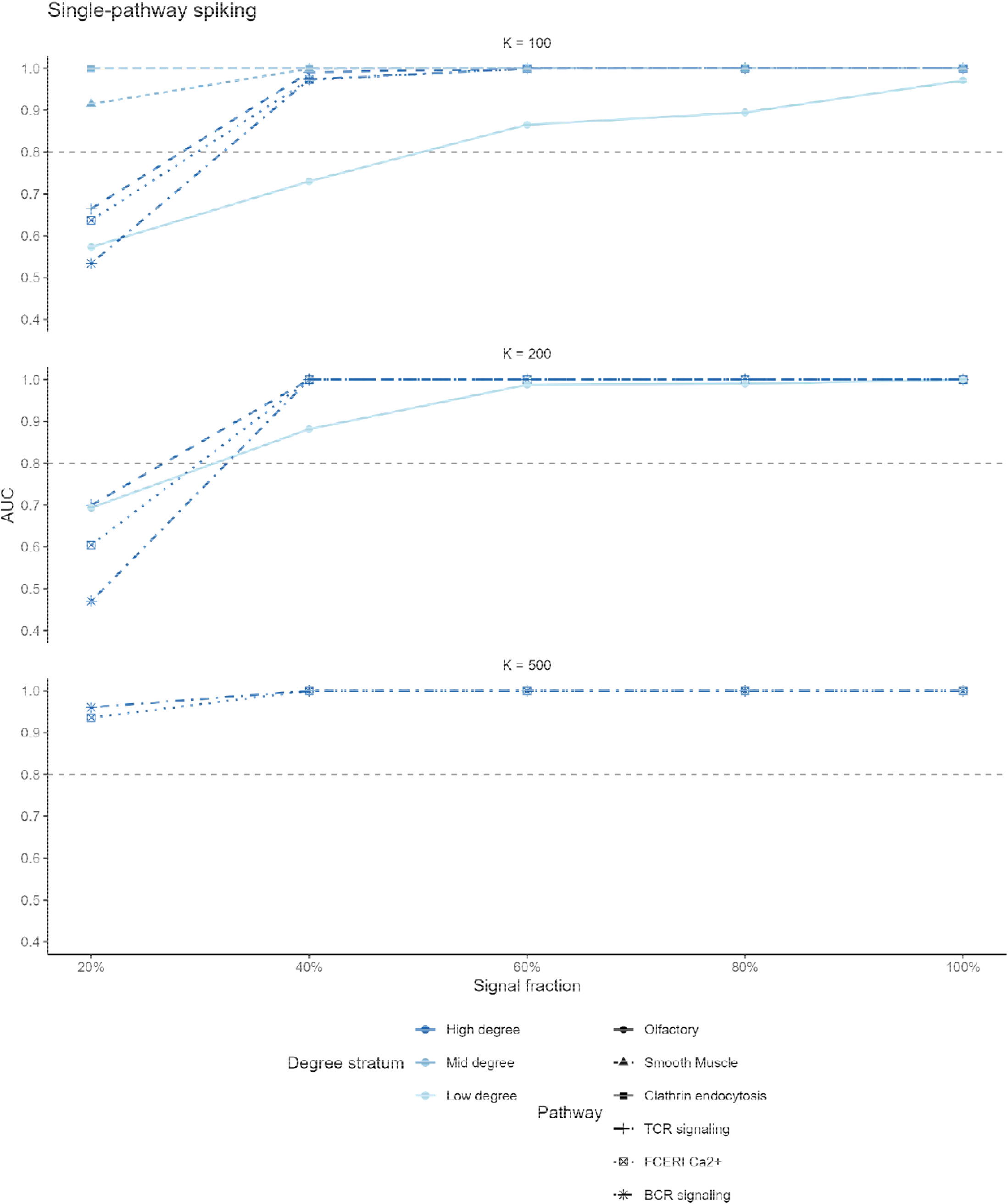
Single-pathway spiking simulations. Single-pathway spiking simulations. AUC of 10-bin Global Moran’s I as a function of signal fraction (proportion of the gene set composed of real pathway genes) for eight individual pathways, faceted by total set size K. Pathways are colored by mean network degree: high (> 150; BCR, FCERI Ca2+, TCR), mid (100–150; Clathrin, Smooth Muscle), and low (< 50; Olfactory). Network-wide mean degree is 34.5. High-degree pathways maintain AUC near 1.0 even at low signal fractions; the low-degree Olfactory pathway degrades earlier. Dashed line marks AUC = 0.80.

In the multi-pathway tier, the same pattern held across degree profiles. High-degree pathway sets (Sets 1–4) achieved AUC ≥ 0.98 at 20% signal fraction and K = 200. Mid-degree Set 5 (Chromatids / Rho / Inflammation) reached AUC ≈ 0.94 under the same conditions. Low-degree Set 6 (Olfactory / Neddylation / mTOR-AMPK) was the hardest case, with AUC ≈ 0.77 at K = 200 and 20% signal, reflecting the genuinely sparse connectivity of those pathway genes. Across all seven pathway sets at K = 200 and 20% signal, the mean AUC under degree-matched nulls remained high (> 0.85 for bins ≤ 5), and the power cost of degree control relative to uniform nulls was consistently small across pathway sets, confirming that degree control imposes minimal power cost even on real pathway clustering (Figure 5). Full pathway-spiking results are in Supplementary Tables S5–S6.

**Figure 5.**
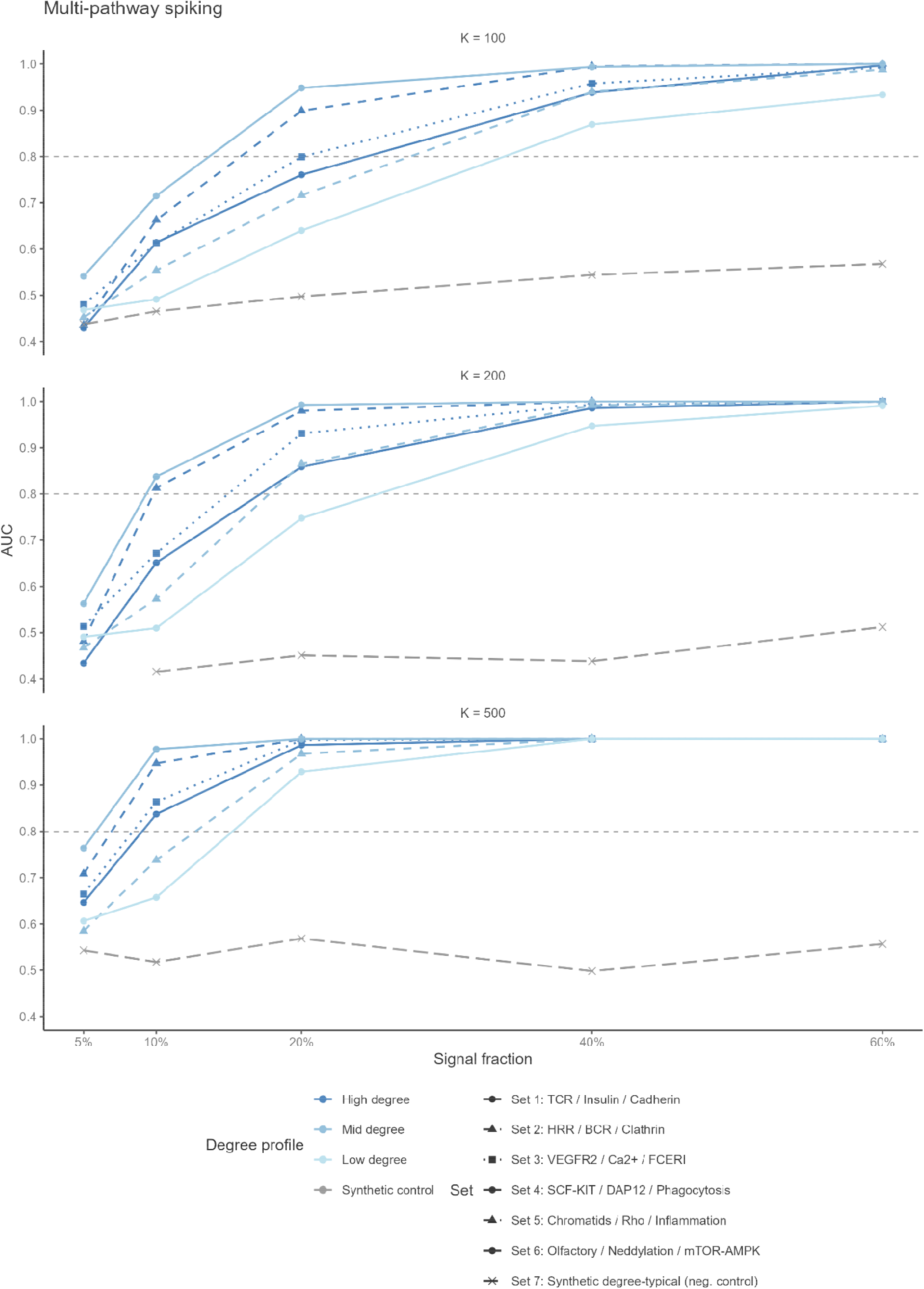
Multi-pathway spiking simulations. AUC of 10-bin Global Moran’s I as a function of signal fraction for seven pathway combinations, faceted by total set size K. Sets are colored by mean network degree of the combined pathway genes: high (> 100; Sets 1–4), mid (Set 5), and low (Set 6). Set 7 is a synthetic degree-typical negative control with no real pathway clustering (expected AUC ≈ 0.5). Dashed line marks AUC = 0.80.

### FIGI colorectal cancer GWAS: a dominant cross-pathway module

We applied MANGO to 115 network-mapped genes from the Functionally Informed GWAS Integration (FIGI) consortium colorectal cancer GWAS. Before running the test, we note a feature of this gene set that matters for interpretation: the FIGI genes are degree-typical, not hub-enriched. Their mean degree in the integrated network is 28.4 against a network-wide mean of 34.5, and a Kolmogorov-Smirnov test against the background degree distribution gives p = 0.83. In other words, this is not a case where the worry is inflated hub-driven apparent clustering, but MANGO’s degree-matched null still applies as principled calibration.

Under MANGO Step 1, the observed Global Moran’s I was 0.0119, with null mean ≈ −2 × 10^−4^ and null SD ≈ 0.0023 (10 bins, 1,000 permutations). The empirical one-sided p-value is 0/1000, i.e. p < 0.001 at this permutation budget: no draw from the degree-matched null matched or exceeded the observed statistic. The p-value is stable across bin choices: p < 0.001 at every setting from 1 to 10 bins. Join Count (the number of within-set edges in the induced subgraph, an adjacency-based complement to Moran’s I) also rejects the null at p = 0.000 at 10 bins.

The induced subgraph decomposed into 82 connected components (Figure 6). Most were singletons (n = 78), together with one 24-gene component, one 8-gene, one 3-gene, and one 2-gene component. The component-level jackknife, which drops each component’s genes from the FIGI set and reruns Step 1, localizes the signal sharply. Removing the 24-gene component raises the Global Moran’s I p-value from < 0.001 to 0.271, i.e. the test is no longer significant without it. Removing any other component, singleton or non-trivial, leaves the p-value at < 0.001. Essentially, the entire Step 1 signal lives in this one module. The reader should note that singletons contribute mechanically little to a global spatial autocorrelation average (they have no within-set neighbors), so the singletons’ null contribution is unsurprising; the informative negative result is that the 8-gene, 3-gene, and 2-gene components also carry no detectable signal.

**Figure 6.**
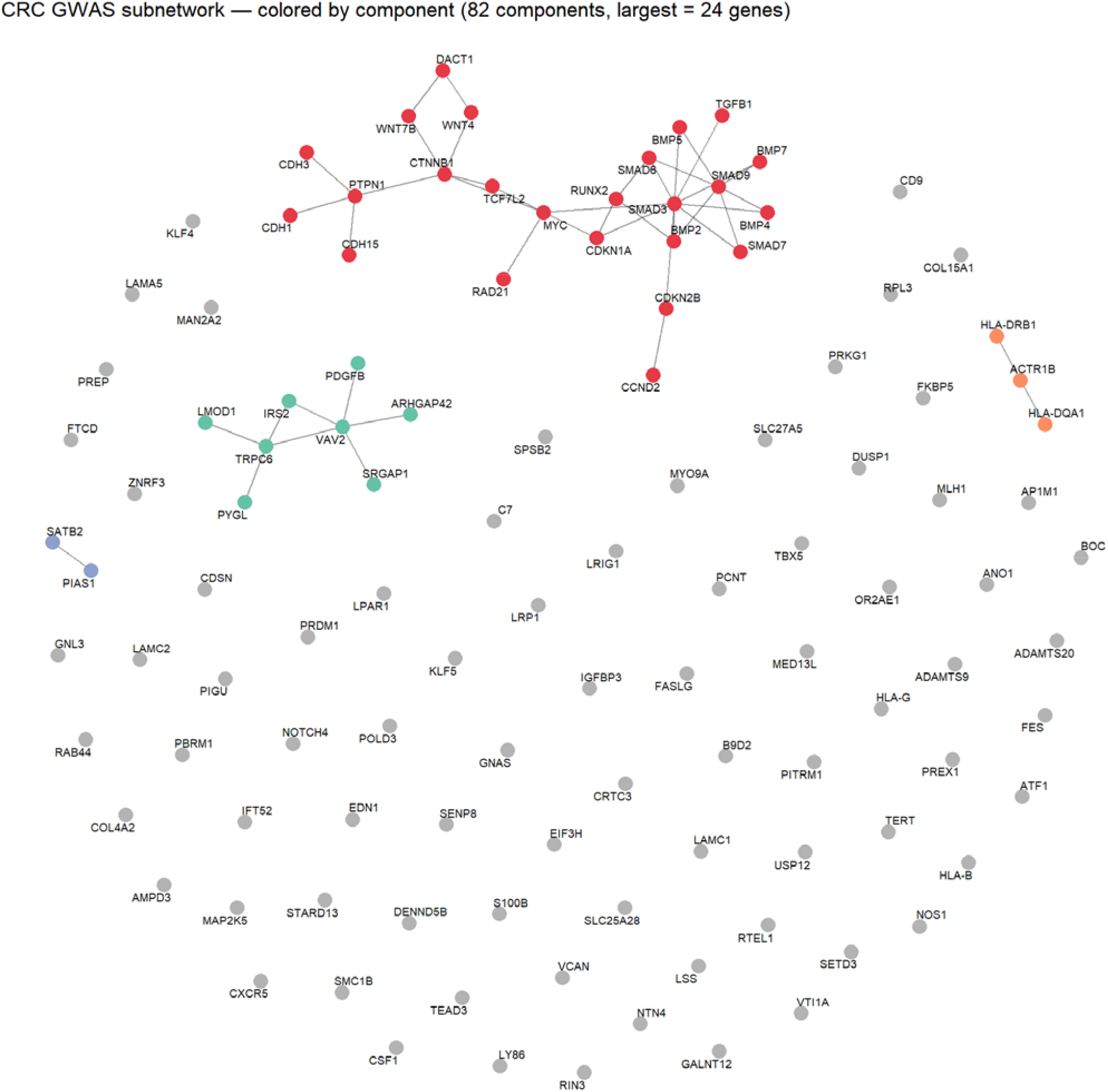
FIGI CRC induced subgraph. The connected components are highlighted. Singletons are in gray.

The dominant 24-gene component spans multiple classical CRC pathways that ORA would return as separate pathway hits. Its genes draw from TGF-β signaling (SMAD3, SMAD6, SMAD7, SMAD9, BMP2, BMP4, BMP5, BMP7, TGFB1), Wnt / cadherin / β-catenin signaling (CTNNB1, MYC, TCF7L2, CDH1, WNT4, WNT7B), and related growth-control and adhesion machinery (CDKN1A, CDKN2B, RUNX2, RAD21, DACT1). Articulation-point analysis within this dominant module identifies four genes whose removal fractures it into two or more pieces: SMAD3 (Δ_components = 2), MYC (+2), CTNNB1 (+2), and PTPN1 (+3). These four bottleneck genes recover the expected architecture of CRC susceptibility: TGF-β/SMAD and Wnt/β-catenin crosstalk routed through MYC, with PTPN1 as a less-appreciated but well-documented CRC bottleneck (overexpressed with a graded increase from normal to dysplasia to carcinoma, inversely associated with survival). SMAD3, MYC, and CTNNB1 are canonical CRC drivers. Four additional articulation points exist in the subgraph (VAV2 and TRPC6 in the 8-gene component; ACTR1B and CDKN2B with Δ = 1), but since the 8-gene component carries no signal under component jackknife, and Δ = 1 corresponds to pruning a leaf rather than fracturing a module, we do not feature these in the main structural narrative (Figure 7).

**Figure 7.**
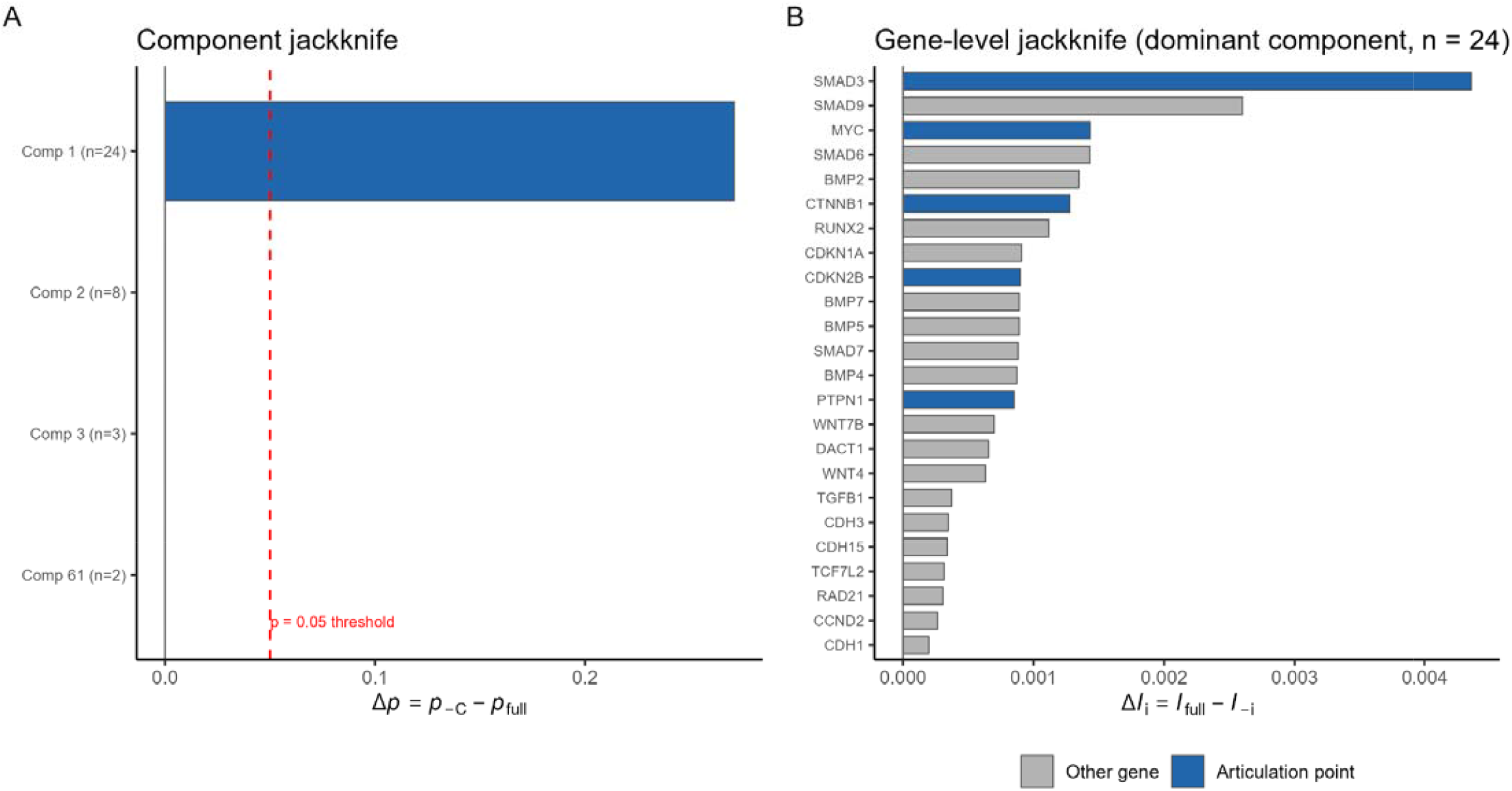
CRC signal attribution. (a) Component jackknife: for each non-trivial component, showing that the dominant component carries the entire signal. (b) Gene-level jackknife within the dominant component, ranked by

Gene-level jackknife tells a consistent story with the component-level result. No single gene’s removal brings the Global Moran’s I test to non-significance: SMAD3 is the largest single-gene contributor, and dropping it yields p = 0.003 (still significant); all other genes give Δp ≤ 0.001. Spearman correlation between a gene’s subgraph degree and its jackknife Δp (for non-singletons only) is ρ = 0.24, p = 0.15 (not significant). The signal is genuinely distributed across the dominant module rather than concentrated in any one driver, which is the expected profile for GWAS gene sets on a real biological network.

### Comparison with existing methods

Table 1 places MANGO alongside representative existing methods (networkGWAS, dmGWAS, HotNet2, LEAN, DOMINO, and the non-network baselines MAGMA and PASCAL) across required inputs, primary test statistic, null model, degree-bias handling, and output granularity. The combination of a Global Moran’s I statistic, a degree-stratified permutation null that preserves network topology, and two-level signal attribution through component and gene-level jackknife is not offered by existing methods in a single framework.

## Discussion

ORA can’t see cross-pathway biology, and network methods that were supposed to fix this inherited a hub-bias confound from curated networks. MANGO addresses both: it holds the network fixed, conditions on the gene set’s connectivity profile, and returns a calibrated p-value for clustering that degree alone can’t explain. The simulations establish that the failure mode of degree-naive tests is not subtle (uniform nulls yield FPR = 1.0 on hub-enriched gene sets), that degree-stratified permutation eliminates it, and that the power cost is negligible. The pathway-spiking results confirm that MANGO detects real biological clustering, not just synthetic BFS geometry.

MANGO sits in a specific place relative to existing methods. The closest comparator is networkGWAS, which also uses a degree-preserving network permutation, but does so in combination with circular SNP permutation inside a mixed-model SNP-set testing framework, and needs individual-level genotype data. The two methods are complementary rather than competing: networkGWAS operates upstream at the SNP-to-gene level, and MANGO operates downstream on the resulting gene set using summary-level inputs. Against NAGA, dmGWAS, HotNet2, and LEAN, MANGO’s contribution is explicit degree control combined with a calibrated p-value, where those methods either skip degree correction or absorb it implicitly into network-propagation parameter tuning. DOMINO’s demonstration that active-module identification methods report GO-enriched modules even on permuted input data is evidence for the problem MANGO solves, not a competing approach; the confounds are different (AMI non-specificity vs. degree-driven clustering inflation) and so are the solutions (per-GO-term empirical calibration vs. degree-stratified permutation). Against MAGMA and PASCAL, which are non-network baselines, MANGO adds information by testing for cross-pathway clustering that predefined pathway boundaries can’t capture.

The closest theoretical precedent is Arthur’s framework for Moran-type statistics on complex networks. Arthur proposes two null models: a data-permutation null (fix edges, permute node values uniformly) and a configuration-model null (fix node values, rewire edges preserving degree). The configuration-model variant is mathematically elegant but biologically destructive for gene-set analysis: it treats the interaction network as a variable to be randomized rather than as a fixed biological constraint. The uniform data-permutation variant preserves the network but does not correct for hub bias. MANGO takes a third option: keep the network fixed but permute gene labels within empirical degree bins rather than uniformly. This is also the positive specification of what the Gillis-Pavlidis critique of guilt-by-association^19^ analysis demands: construct the null conditional on degree, so that significance certifies clustering beyond what degree alone would produce.

The CRC application illustrates what MANGO delivers that ORA does not. Classical ORA on the FIGI gene set would return separate significant hits across TGF-β, Alzheimer’s, Gonadotropin, and cadherin signaling pathways, as shown in Guaderman’s^29^. The reader is left to synthesize those hits narratively. MANGO replaces that synthesis with a single statistical object: one autocorrelated 24-gene module spanning all four pathways, whose contribution to the global signal is directly verifiable (dropping it raises the p-value from < 0.001 to 0.271, while dropping any other component leaves it unchanged). Gene-level jackknife identifies SMAD3, MYC, CTNNB1, and PTPN1 as the top statistical drivers within this module, all of which map onto established CRC biology, which we treat as a consistency demonstration rather than independent discovery.

Several limitations merit discussion. As with any test of spatial autocorrelation, detection power depends on both gene-set size and signal fraction: a small gene set with a low proportion of genuinely clustered genes presents a weak signal that any method would struggle to detect. The pathway-spiking simulations quantify this tradeoff across K = 100, 200, and 500, and show that MANGO achieves AUC > 0.80 at 20% signal fraction for most pathway degree profiles at K = 200, but low-degree pathways require higher signal fractions or larger sets. MANGO does not remove degree bias from curated interaction networks; it controls for degree so that any clustering signal that survives the degree-matched null cannot be explained by the degree distribution of the gene set alone, whether that degree reflects biological centrality, literature attention, or both. An under-annotated gene with real biological importance may simply not appear in the induced subgraph. The current framework treats all edges as binary and unweighted. Extension to weighted networks with principled edge-confidence scores is conceptually aligned with the existing construction. The present study uses a single integrated network (Reactome, PANTHER, KEGG via PathwayCommons) and a single disease application. Ongoing analyses in schizophrenia, Type 2 diabetes, and breast cancer will test generalizability, and sensitivity analyses across GO-CAM, STRING, BioGRID, and HumanNet will quantify dependence on the specific curation choices.

## Conclusion

MANGO is a topology-preserving statistical framework for gene-set analysis built on spatial autocorrelation. It separates study-driven hub bias from biological signal by calibrating Global Moran’s I against a degree-matched permutation null on a fixed interaction network, and attributes the resulting signal to specific modules through component jackknife and to individual driver genes through gene-level jackknife.

Three findings support the framework. First, degree-stratified permutation eliminates the false-positive inflation that uniform nulls produce on hub-enriched gene sets, with negligible power cost on degree-typical signals. Second, pathway-spiking simulations confirm that MANGO detects real biological clustering across a range of pathway sizes and degree profiles, not just synthetic scenarios. Third, the CRC application demonstrates end-to-end utility: a single 24-gene module spanning TGF-β, Wnt/cadherin, and related pathways accounts for the entire global signal, with gene-level attribution identifying known CRC drivers.

## References

1. García-Campos, M. A., Espinal-Enríquez, J. & Hernández-Lemus, E. Pathway Analysis: State of the Art. Front. Physiol. 6, 383 (2015).

2. Holmans, P. Statistical methods for pathway analysis of genome-wide data for association with complex genetic traits. Adv. Genet. 72, 141–179 (2010).

3. Milacic, M. et al. The Reactome Pathway Knowledgebase 2024. Nucleic Acids Res. 52, D672–D678 (2024).

4. Kanehisa, M., Furumichi, M., Sato, Y., Matsuura, Y. & Ishiguro-Watanabe, M. KEGG: biological systems database as a model of the real world. Nucleic Acids Res. 53, D672–D677 (2025).

5. Thomas, P. D. et al. PANTHER: Making genome-scale phylogenetics accessible to all. Protein Sci. 31, 8–22 (2022).

6. Ashburner, M. et al. Gene Ontology: tool for the unification of biology. Nat. Genet. 25, 25–29 (2000).

7. Consortium, T. G. O. et al. The Gene Ontology knowledgebase in 2026. 10.1093/nar/gkaf1292.

8. Khatri, P., Sirota, M. & Butte, A. J. Ten Years of Pathway Analysis: Current Approaches and Outstanding Challenges. PLOS Comput. Biol. 8, e1002375 (2012).

9. Ziemann, M., Schroeter, B. & Bora, A. Two subtle problems with overrepresentation analysis. Bioinforma. Adv. 4, vbae159 (2024).

10. de Oliveira, F. H. S., Gomes, F. A. & Feltes, B. C. Benchmarking multiple gene ontology enrichment tools reveals high biological significance, ranking, and stringency heterogeneity among datasets. Front. Bioinforma. 6, (2026).

11. Menche, J. et al. Uncovering disease-disease relationships through the incomplete human interactome. Science 347, 1257601 (2015).

12. Leiserson, M. D. M. et al. Pan-cancer network analysis identifies combinations of rare somatic mutations across pathways and protein complexes. Nat. Genet. 47, 106–114 (2015).

13. Jia, P., Zheng, S., Long, J., Zheng, W. & Zhao, Z. dmGWAS: dense module searching for genome-wide association studies in protein–protein interaction networks. 10.1093/bioinformatics/btq615.

14. Levi, H., Rahmanian, N., Elkon, R. & Shamir, R. The DOMINO web-server for active module identification analysis. 10.1093/bioinformatics/btac067.

15. Gwinner, F. et al. Network-based analysis of omics data: the LEAN method. 10.1093/bioinformatics/btw676.

16. Muzio, G. et al. networkGWAS: a network-based approach to discover genetic associations. Bioinformatics 39, btad370 (2023).

17. Blumenthal, D. B. et al. Emergence of power law distributions in protein-protein interaction networks through study bias. eLife 13, e99951 (2024).

18. Schaefer, M. H., Serrano, L. & Andrade-Navarro, M. A. Correcting for the study bias associated with protein–protein interaction measurements reveals differences between protein degree distributions from different cancer types. Front. Genet. 6, (2015).

19. Gillis, J. & Pavlidis, P. The Impact of Multifunctional Genes on ‘Guilt by Association’ Analysis. PLOS ONE 6, e17258 (2011).

20. Levi, H., Elkon, R. & Shamir, R. DOMINO: a networkLbased active module identification algorithm with reduced rate of false calls. Mol. Syst. Biol. 17, MSB20209593 (2021).

21. Maslov, S. & Sneppen, K. Specificity and Stability in Topology of Protein Networks. Science 296, 910–913 (2002).

22. Moran, P. a. P. NOTES ON CONTINUOUS STOCHASTIC PHENOMENA. 10.1093/biomet/37.1-2.17.

23. Getis, A. Cliff, A.D. and Ord, J.K. 1973: Spatial autocorrelation. London: Pion. Prog. Hum. Geogr. 19, 245–249 (1995).

24. Arthur, R. Correlation and autocorrelation of data on complex networks. EPJ Data Sci. 14, 6 (2025).

25. Palla, G. et al. Squidpy: a scalable framework for spatial omics analysis. Nat. Methods 19, 171–178 (2022).

26. Rodchenkov, I. et al. Pathway Commons 2019 Update: integration, analysis and exploration of pathway data. Nucleic Acids Res. 48, D489–D497 (2020).

27. Demir, E. et al. The BioPAX community standard for pathway data sharing. Nat. Biotechnol. 28, 935–942 (2010).

28. Latora, V. & Marchiori, M. Efficient Behavior of Small-World Networks. Phys. Rev. Lett. 87, 198701 (2001).

29. Gauderman, W. J. et al. Pathway polygenic risk scores (pPRS) for the analysis of gene-environment interaction. PLOS Genet. 21, e1011543 (2025).

